# Activation of NLRP3 inflammasome/pyroptosis by protease inhibitor atazanavir at high concentrations is mediated through mitochondrial dysfunction

**DOI:** 10.1101/2024.06.18.599585

**Authors:** Lei Zhou, Moises Cosme, Min Li, Velma Eduafo, Dereck Amakye Jnr, Elijah Paintsil

**Affiliations:** Department of Pediatrics, Yale School of Medicine, New Haven, Connecticut, USA; Department of Pharmacology, Yale School of Medicine, New Haven, Connecticut, USA; Department of Epidemiology & Public Health, Yale School of Medicine, New Haven, Connecticut, USA

## Abstract

Protease inhibitor (PI)-based antiretroviral therapy increases CD4 and CD8 cell counts through anti-apoptosis mechanism. However, there are emerging reports that PIs could have pro-apoptotic effects; thus, PIs may have biphasic effect on apoptosis. We hypothesized that PI-induced apoptosis may be mediated through PI-induced mitochondrial dysfunction resulting in increased production of reactive oxygen species (ROS).

To test this hypothesis, we used human T lymphoblastoid cell line (CEM) cultured with increasing concentrations of atazanavir (ATV). We assessed mitochondrial function—i.e., cell growth, apoptosis, mitochondrial membrane potential (ΔΨ), ROS, and electron transport chain (ETC) proteins. Apoptosis pathway genes were interrogated using the Human Apoptosis RT² Profiler PCR Array kit (QIAGEN), followed by quantitative PCR for validation.

CEM cells treated with 15, 22.5, and 30 µM of ATV resulted in significant reduction in cell growth and increased apoptosis. Further, high concentrations of ATV resulted in decreased mitochondrial ΔΨ, increased ROS production, and decreased protein expression of ETC I, II, III, and V. The following apoptosis pathway genes—caspase-1, BCL2A1, TP73 and TNFRSF1B—were differentially expressed. Caspase-1 is known to play a role in inducing NLRP3 inflammasome/pyroptosis pathway. We validated this with qPCR of the genes in the NLRP3 inflammasome pathway. Of note, NLRP3 inhibitor MCC950 and caspase-1 inhibitor YVAD reversed mitochondrial dysfunction and cell death. Our findings suggest that ATV-induced cell death is through the NLRP3 inflammasome/pyroptosis pathway. ATV-induced mitochondrial dysfunction plays an important role in the regulation of ATV-activated NLRP3 inflammasome pathway.

## Introduction

HIV protease inhibitors (PIs) target the last step of the replication cycle of HIV. They inhibit HIV protease from cleaving the polyproteins, thus producing immature, noninfectious viral particles [1]. The advent of PI-based antiretroviral therapy (ART) regimens resulted in a decrease in AIDS-related morbidity and mortality [2–4]. In addition to inhibition of HIV protease resulting in decrease viral replication, PIs also modulate apoptosis [5–7]. Several studies have reported that PIs induce a decrease in apoptosis of lymphocytes leading to increase in CD4 and CD8 cell numbers [7–9]. However, other studies have reported that PIs have pro-apoptotic effects, particularly at higher concentrations [7, 10–13]. High concentrations of ritonavir ((10-50 µM) increased apoptosis in murine and human tumor cell lines; the concentrations of ritonavir used in these experiments were over 15 times the serum concentration achieved in adults on ritonavir treatment [10]. Moreover, when primary cells and cell lines of Adult T-cell Leukemia were incubated with 20-40 µM of ritonavir, there was a five-fold increase in apoptosis [11]. Lopinavir and ritonavir treatment also induced apoptosis in neuroblastoma cell line at 25 and 50 µM concentrations [12]. Treatment of monocytes and CD4 T cells with 10 μM indinavir or saquinavir significantly increased apoptosis [7]. Nelfinavir was reported to induce apoptosis in medullary thyroid cancer cells at therapeutic serum concentration (10 µM) [13]. Atazanavir (ATV), the most popular PI, has favorable safety profile, but its effect on apoptosis has not been well-characterized.

Other classes of antiretroviral drugs have been associated with increasing apoptosis; e.g., nucleoside reverse transcriptase inhibitors (NRTIs) and non-nucleoside reverse transcriptase inhibitors (NNRTIs). 2′, 3′-dideoxycytidine (ddC, an NRTI) induced apoptosis and neuronal death, with oxidative stress and mitochondrial dysfunction proposed as key mechanisms of toxicity [14]. Efavirenz (EFV, an NNRTI) treatment induced caspase- and mitochondrion-dependent apoptosis of Jurkat T cells and human peripheral blood mononuclear cells[15]. This is consistent with our previous report that EFV induced mitochondria dysfunction and apoptosis in human T lymphoblastoid cells [16]. Even though PIs do not inhibit Pol-γ, PIs have been associated with increased ROS production, decreased mitochondrial ΔΨ, and apoptosis [17]; these are functions of the mitochondria.

In general, apoptosis refers to programmed cell death. However, programmed cell death can be categorized into different cell death modalities—pyroptosis, apoptosis, autophagy, and ferroptosis. Pyroptosis is an inflammation-related programmed cell death that is triggered by various pathological stimuli and plays a critical role in innate immunity against bacterial and viral infections [18, 19]. The type of programmed cell death caused by PIs is not well-known. We hypothesized that mitochondrial dysfunction leading to increased ROS production will be in the causal pathway of PI-induced apoptosis. We cultured human T lymphoblastoid cell line (CEM cells) with increasing concentrations of ATV to investigate its effect on cell death and underlying mechanism(s).

## MATERIALS & METHODS

### Cell culture and drugs

CEM cells were cultured in RPMI 1640 supplemented with 10% dialyzed fetal calf serum (Thermo Fisher Scientific, NY, USA) and incubated at 37°C in a 5% CO2 humidified environment. ATV was from the NIH HIV Reagent Program. The maximum concentration (Cmax) is the highest concentration of a drug in the blood, cerebrospinal fluid, or target organ after a dose is given. The Cmax of ATV in plasma after a standard dose ranges from 1.5 to 8 µM in patients [20–22]. 4 × 10^4^/ml cells (total volume of 30 ml) were cultured with different concentrations of ATV, multiplicities of Cmax— 0, 2.5, 5.0, 7.5, 15, 22.5, and 30.0 µM. The cells were passaged every 4 days and fresh media containing drug was added. We harvested the cells at days 2, 4, and 8 for cell count and the experiments described below. For experiments involving caspase-1 and NLRP3 inhibitors, CEM cells were pre-incubated with either 20 µM of AC-YVAD-AMK or 1 µM of MCC950 for 1h, followed by treatment with dimethyl sulfoxide (DMSO) and 30 µM ATV. Cell harvesting was conducted after 6 hours on the same day. Aliquots of cells were stained with Trypan blue to distinguish live cell from dead cells and the number of live cells was counted using hemocytometer. For each treatment condition, at least three cell culture experiments were conducted on different occasions.

### Flow cytometry analyses

Cells treated with DMSO and different concentrations of ATV were harvested at days 2, 4, and 8. Cells from cell culture suspension were pelleted and the media was aspirated before washing cells with cold phosphate buffered saline (PBS). Cells were stained following manufacturer’s instructions of detection kits as previously published [23]. Apoptosis was assessed by staining with an FITC Annexin V Apoptosis Detection Kit I (BD, San Jose, CA), and early apoptotic cells (Annexin V+/PI-) or late apoptotic cells (Annexin V+/PI+) were evaluated by double staining with annexin V–FITC and PI for 15 min.

Mitochondrial ΔΨ was analyzed using TMRE-ΔΨ assay Kit (Abcam, Cambridge, MA). TMRE (tetramethylrhodamine, ethyl ester) is a cell permeant, positive-charged, red-orange dye that readily accumulates in active mitochondria due to their relative negative charge. Depolarized or inactive mitochondria have decreased membrane potential and fail to sequester TMRE. Cells were stained with 200 nM TMRE respectively for 20 min at room temperature.

Production of ROS was determined using MitoSOX™ Red mitochondrial superoxide indicator. Cells were incubated with 5 μM MitoSOX™ reagent working solution at room temperature for 10 min. After incubation with corresponding dyes, reactions were stopped by adding 300 µl of PBS and immediately analyzed on BD LSR II flow cytometer (BD, San Jose, CA) and with BD FACS Diva software. Each experiment was done in triplicates and repeated on at least three different occasions.

### RT^2^ Profiler PCR Arrays

RNA was extracted from aliquots of cells harvested at day 8 treated with different concentrations of ATV using RNeasy Plus Mini Kit (Qiagen, Germantown, MD) according to the manufacturer’s instructions. QuantiTect Reverse Transcription kit (Qiagen, Germantown, MD) was used to reverse transcribe RNA to cDNA. To identify genes of interest, a 96 well RT^2^ profiler PCR array human apoptosis kit (Qiagen, Germantown, MD) containing 84 apoptotic genes was used according to manufacturer’s instructions. Real-time quantitative PCR was performed in triplicate with a final volume of 25 µL per well consisting of 12.5 ng cDNA, 2x SYBR Green (Thermo Fisher Scientific, Waltham, MA), 10 µM forward primer, and 10 µM reverse primer. Thermal cycling was performed using iCycler iQ5 (Bio-Rad) as follows: 95.0°C for 10 minutes, 40 cycles of 95°C for 15 seconds and 60° C for 1 minute, during which real time qPCR data was collected, and 91 cycles lasting 6 seconds, starting at 50.0° C and increasing in increments of 0.5°C per cycle to retrieve the melting curve. Data was then analyzed using iQ5 Optical System Software (Bio-Rad Laboratories v 2.1).

### Quantitative real-time polymerase chain reaction

RNA was extracted from aliquots of cells harvested at day 8 treated with different concentrations of ATV using RNeasy Plus Mini Kit (Qiagen, MD, USA) according to the manufacturer’s instructions and used in quantitative real-time PCR as previously described [23]. The genes of interest and their sequences of primers are listed in Table 1. Glyceraldehyde 3-phosphate dehydrogenase (GAPDH) gene was used as an internal control for all reactions. The threshold cycle (CT) values of the genes were determined for each treatment condition. The fold-change in gene expression was calculated as 2^-ΔΔCT^; where ΔΔCT = ΔCT(treated) – ΔCT(control); ΔCT(treated) = CT(gene of interest) – CT(GAPDH); ΔCT(control)= CT(gene of interest) – CT(GAPDH). All of the samples were run in duplicate in at least three independent experiments.

**Table 1.**
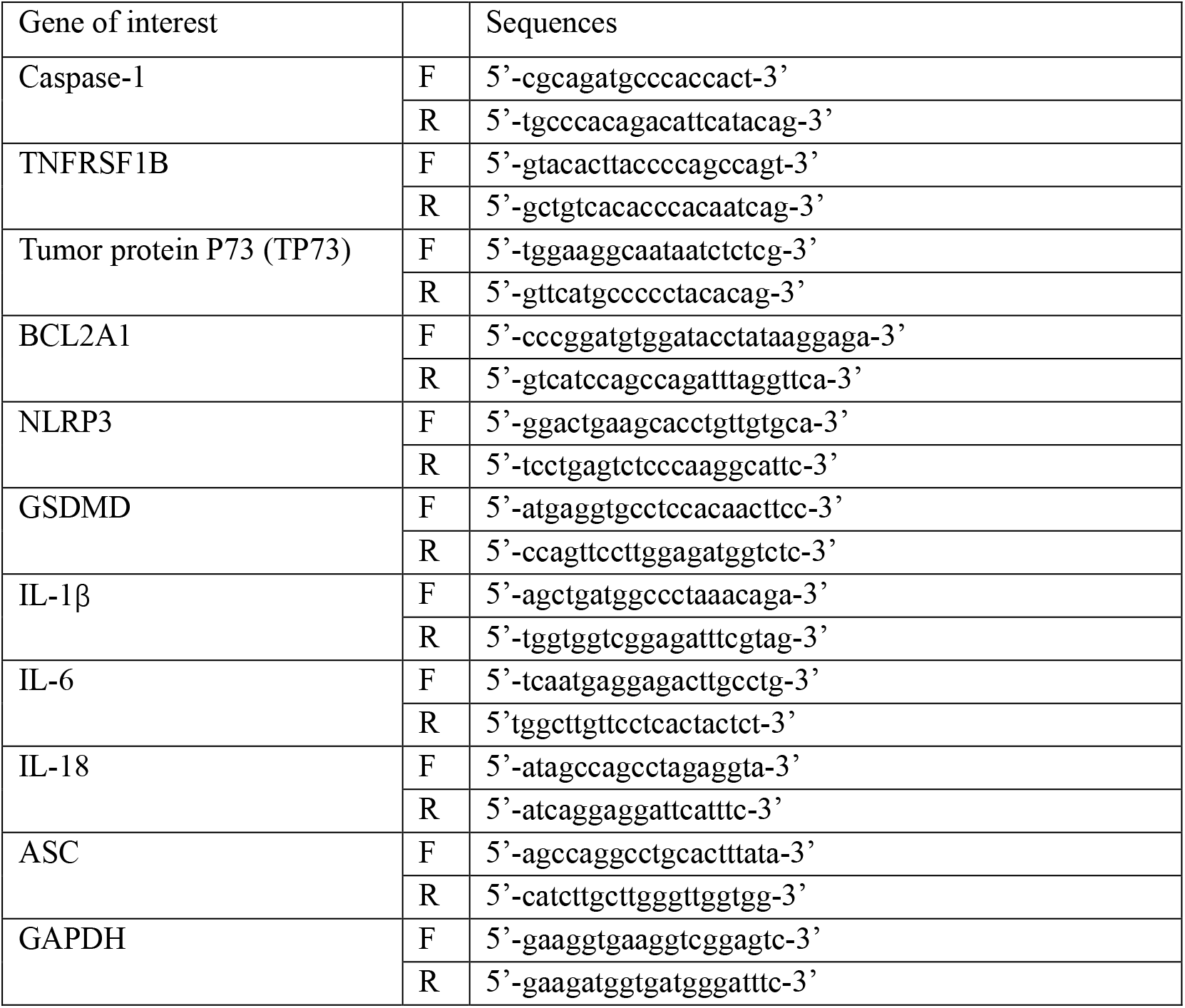
Sequences of qPCR primers.

### Protein extraction and western blot analyses

We extracted total protein from cells using RIPA lysis and extraction buffer (Millipore Sigma. Darmstadt, Germany) according to the manufacturer’s instructions. Extracted proteins were quantified using Pierce™ BCA Protein Assay Kits (ThermoScientific, Rockford, IL) according to the manufacturer’s instructions. Western blot analysis was performed as described previously [23] using total cell protein extracts from CEM cells treated with ATV and DMSO. Tubulin was used as the house keeping gene. Primary antibodies: complex V (CV-APT5A), complex III (CIII-UQCRC2, complex IV (CIV-MTCO1), complex II (CII-SDH8), and complex I (CI-NDUFB8) were used at 1:500 (Abcam, Waltham, MA. USA); pro-CASP1 (dilution 1:500; cat. no. 3866S; Cell Signaling Technology, Danvers, MA); secondary antibodies were HRP conjugated anti-rabbit antibodies at 1:2000 (Cell Signaling Technology, Danvers, MA, USA) and HRP conjugated anti-mouse antibodies: 1:7500 (GE, Pittsburg, PA, USA). Enhanced chemiluminescence substrate was used for signal development (PerkinElmer, Shelton, CT, USA).

### Data and statistical analysis

All statistical analyses were performed with GraphPad Prism software with the Student’s t-test. Data are expressed as means ± SD and significance was achieved when p value was <0.05.

## Results

### High concentrations of ATV increased cell death in human T-lymphoblast cells

We treated CEM cells with DMSO or increasing concentrations of ATV (0-30 µM) and monitored for cell growth. ATV treatment concentrations of 15, 22.5, and 30 µM resulted in significant reduction in cell growth compared to DMSO treated cells (Figure 1A). CEM cells treated with increasing concentration of ATV displayed significant increase in the amount of cell death starting at day 2 at 22.5 µM, with more significant increase at 15, 22.5, and 30 µM at days 4 and 8. At lower concentrations of ATV, around 1× Cmax, no significant increase or decrease in cell death was observed. ATV 30 µM treatment resulted in the greatest amount of cell death compared to all other treatment conditions.

**Figure 1.**
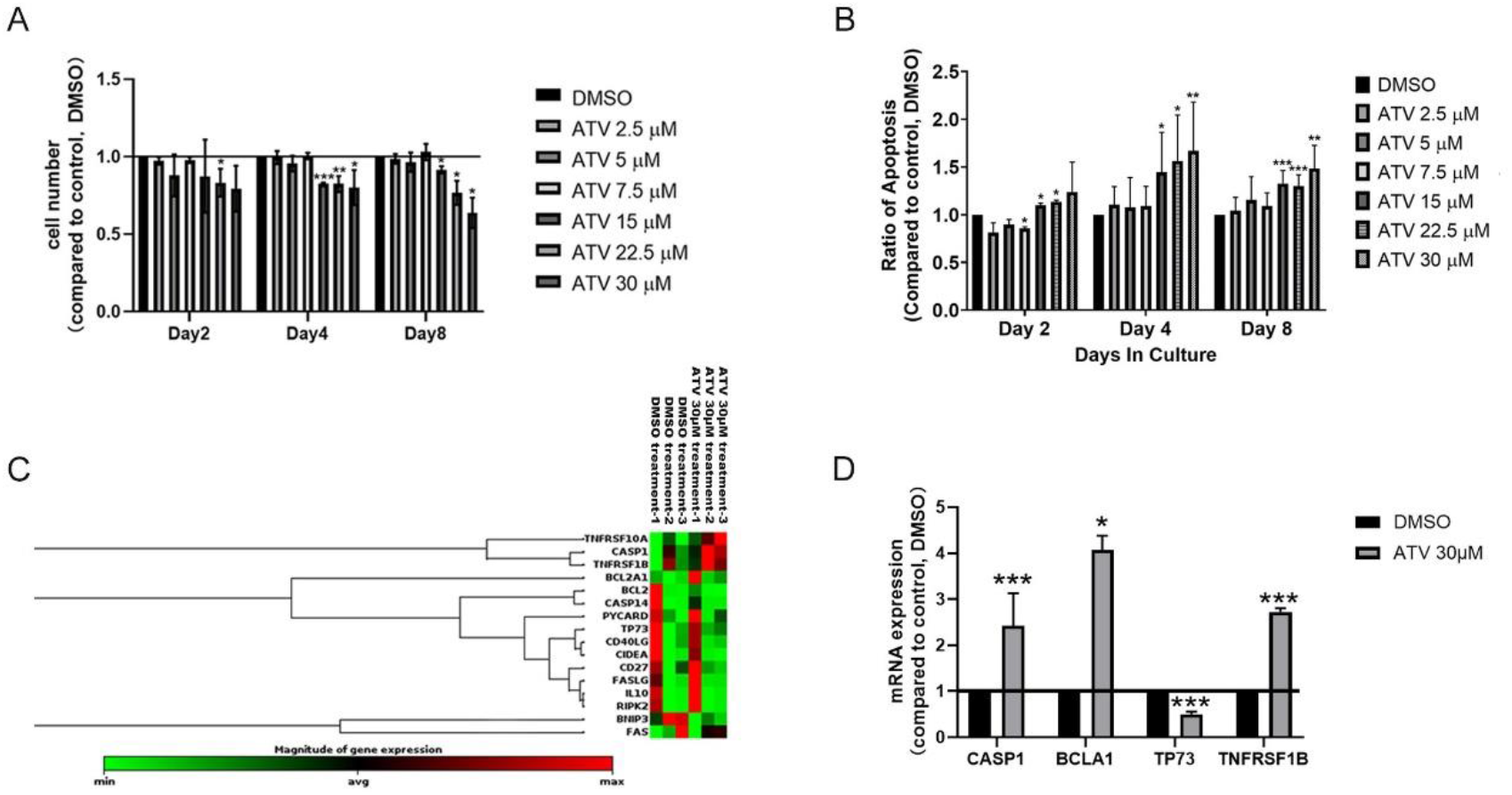
Effect of atazanavir (ATV) on cell viability and apoptosis in CEM cells. A. Effect of ATV on cell viability. CEM cells were cultured with dimethyl sulfoxide (DMSO) and different concentrations of ATV (2.5, 5.0, 7.5, 15, 22.5 and 30.0 µM). At days 2, 4, and 8, aliquots of the cell culture were collected, and live cells were counted with trypan blue staining using hemocytometer. B. Apoptosis was assessed by staining with an FITC Annexin V Apoptosis Detection Kit I. Bar graphs represent proportion of apoptotic cells after 2, 4, and 8 days of treatment compared to DMSO-treated cells. C. Heatmap of RT^2^ Profiler PCR Arrays result. Cells collected under DMSO and 30 µM ATV treatment at day 8 were used for comparison. D. mRNA expression levels of results of caspase-1 (CASP1), BCL2A1, tumor protein p73 (TP73) and TNFRSF1B by qPCR. Primers used are listed in Table 1. Data represent at least 3 independent experiments and plotted as mean ± SD. P-values are two sided and considered significant if <0.05 (*), < 0.01 (**), or <0.001 (***).

We then determined cell death/apoptosis using PI/Annexin-V flow cytometry at days 2, 4 and 8 using previously published protocol (21). As shown in Figure 1B, we observed significantly increased apoptosis in cells treated with 22.5 µM of ATV from day 2 and cells treated with 15 and 30 µM from day 4 compared to cells treated with DMSO.

### High concentrations of ATV increased expression of caspase-1 gene

To investigate the underlying mechanism(s) of ATV-induced cell death, we used RT^2^ profiler PCR array human apoptosis kit containing 84 genes involved in apoptosis pathway. CEM cells treated with DMSO versus 30 µM ATV for 8 days were used. Figure 1C is the heatmap of genes with fold change over 1.5. We observed differential expression levels of caspase-1, BCL2A1, tumor protein p73 (TP73) and TNFRSF1B between DMSO and ATV treatment. We validated these findings with qPCRs. As shown in Figure 1D, 30 µM ATV treatment induced significantly higher expression levels of caspase-1, BCL2A1, TP73, and TNFRSF1B genes than in DMSO treated cells. BCL2A1 is an anti-apoptotic gene, it inhibits apoptosis by preventing pore formation in the mitochondrial outer membrane [24]. It also serves as an early-response gene protecting cells from apoptosis [25]. TP73 participates in the apoptotic response to DNA damage [26]. TNFRSF1B, or TNF-alpha, is a known inflammatory cytokine that may lead to cell signaling events that trigger inflammasome complex formation [27].

Among these genes, caspase-1 drew our attention as it plays fundamental role in inducing pyroptotic cell death.

### High concentrations of ATV activated the NLRP3 inflammasome pathway

We speculated that caspase-1 involved pyroptosis pathway could be the underlying mechanism of ATV-induced cell death. We investigated this using qPCR to determine the expression levels of key genes in NLRP3 inflammasome/pyroptosis pathway. CEM cells were treated with DMSO (control) and 30 µM ATV for 8 days. As shown in Figure 2A, 30 µM ATV significantly enhanced the mRNA expressions of NLRP3, ASC, and GSDMD. The expression level of the effector cytokine IL-1β, IL-18 and IL-6 were also significantly increased. Activation of the NLRP3 inflammasome was also confirmed by a reduction in the protein expression of pro-caspase-1 in ATV treated cells (Figure 2B).

**Figure 2.**
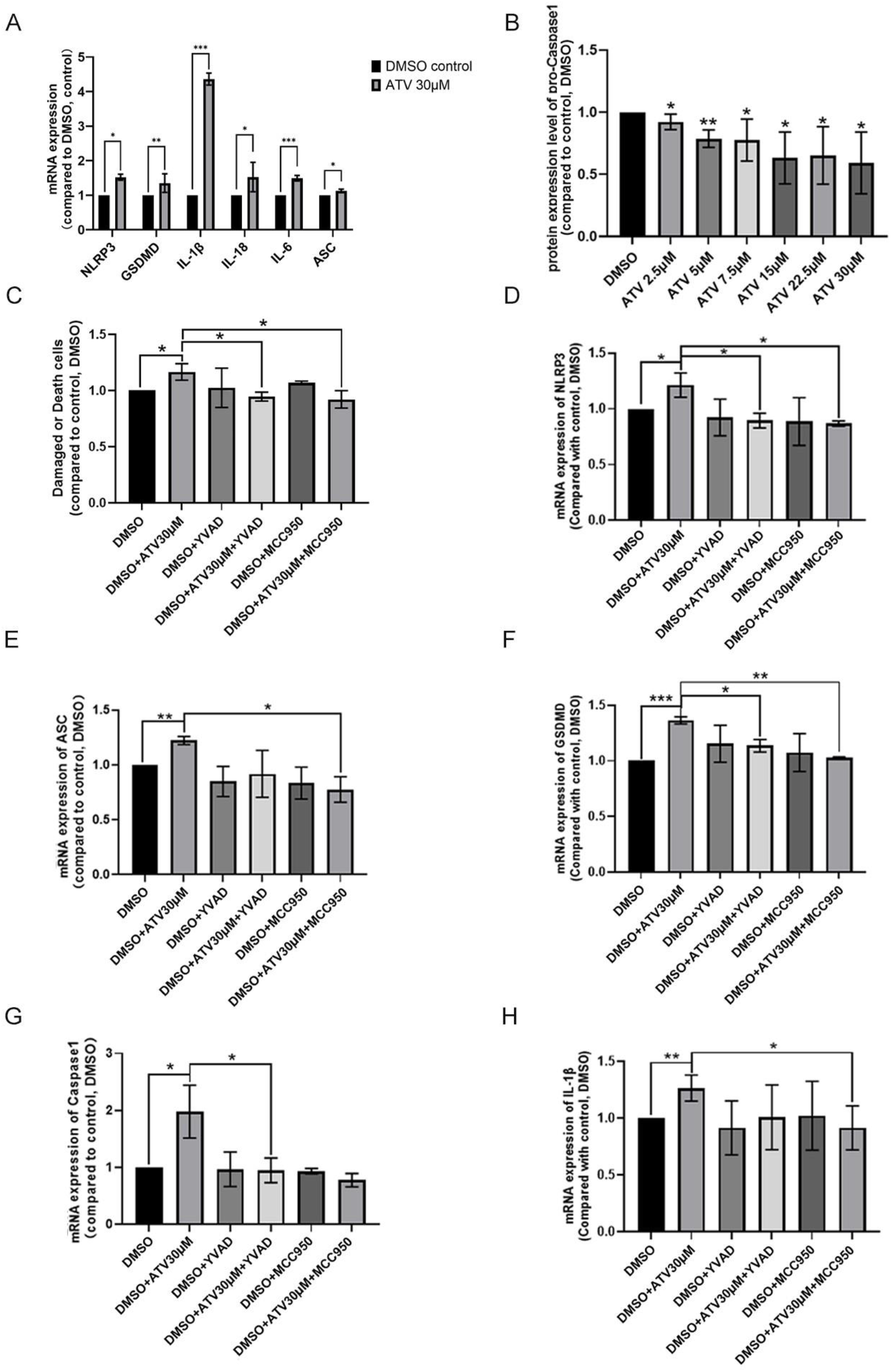
Effect of atazanavir (ATV) on NLRP3 inflammatory pathway. CEM cells were cultured with dimethyl sulfoxide (DMSO) and different concentrations of ATV (2.5, 5.0, 7.5, 15, 22.5 and 30.0 µM) for 8 days. At day 8, the cells were harvested, and RNA and protein were extracted. CEM cells were pre-incubated with AC-YVAD-AMK (20 µM) or MCC950 (1 µM) for 1 h before ATV treatment. A. mRNA expression level of key genes in NLRP3 inflammasome pathway. Real time qPCR of NLRP3, ASC, IL-1β, IL-6, IL-18 were performed. The expression levels of the genes were normalized to the expression of housekeeping gene, GAPDH. Data (mean ± SD, n = 3), are represented as fold change in expression compared with cells treated with DMSO. B. Cells treated for 6 h were harvested, and damage or dead cells was assessed by staining with an FITC Annexin V Apoptosis Detection Kit I. Bar graphs represent proportion of damage or dead cells of ATV treatment compared to DMSO-treated cells. C. Western blot result of pro-caspase-1. The density of the pro-caspase-1 was normalized to that of GAPDH. D-H. cells after treating 6 h were harvested and RNA were extracted. mRNA expression level of NLRP3 (D), ASC (E), GSDMD (F), Caspase-1 (F) and IL-1β (H) by RT-qPCR. Data (mean ±SD) were compared with DMSO treated cells and analyzed by Student’s t-test. P-values are two sided and considered significant if <0.05 (*), < 0.01 (**), or <0.001 (***).

We further used caspase-1 inhibitor AC-YVAD-CMK and NLRP3 inhibitor MCC950 to validate the above results. CEM cells were pre-incubated with 20 µM of AC-YVAD-AMK or 1 µM of MCC950 for 1h and then treated with DMSO or 30 µM ATV for 6 hours. We observed a significant reduction in the number dead cells when cells were pre-incubated with AC-YVAD-CMK or MCC950 before administering ATV treatment compared to treatment with ATV alone (Figure 2C). In addition, pre-incubation of cells with AC-YVAD-CMK reversed the increased mRNA expression of NLRP3, GSDMD and caspase-1 in cells treated with 30 µM ATV (Figure 2D, F and G). Pre-incubation with MCC950 decreased the expressions of NLRP3, ASC and GSDMD, and IL-1β (Figure 2D, E, F and H). These results indicated that AC-YVAD-CMK and MCC950 could rescue human T-lymphoblast cells from ATV-induced pyroptosis.

### High concentrations of ATV caused mitochondrial dysfunction

We further investigated whether ATV treatment caused mitochondrial dysfunction in CEM cells. As reduction of mitochondrial ΔΨ is a key indicator of mitochondrial dysfunction [30], we firstly tested the effect of ATV treatment on mitochondrial ΔΨ of CEM cells using TMRE (tetramethylrhodamine, ethyl ester). Cells treated with 22.5 and 30 µM of ATV showed statistically significant higher proportion of cells with decreased mitochondrial ΔΨ in a concentration-dependent manner (Figure 3A). No statistically significant difference in mitochondrial ΔΨ was observed between cells treated with 1× Cmax of ATV and DMSO.

**Figure 3.**
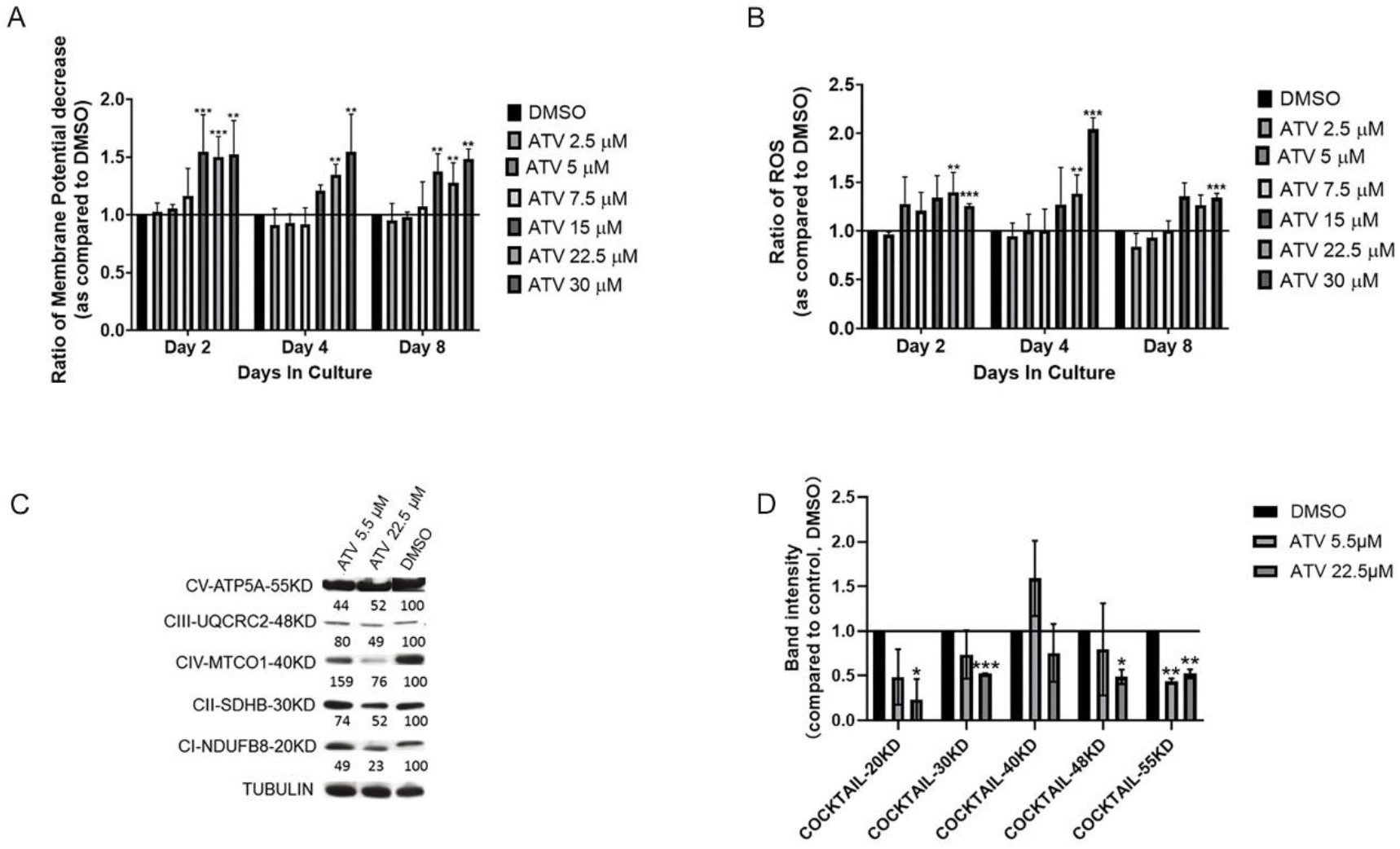
Effect of atazanavir (ATV) treatment on mitochondrial function. CEM cells were cultured with dimethyl sulfoxide (DMSO) and different concentrations of ATV treatment (2.5, 5.0, 7.5, 15, 22.5 and 30.0 µM) for 8 days. A. At days 2, 4, and 8, the cells were harvested and interrogated for mitochondrial membrane potential (ΔΨ) using TMRE-ΔΨ Assay Kit with flow cytometry. The bar graph represents the relative decrease of mitochondrial ΔΨ in cells treated with antiretroviral drugs compared to DMSO-treated control cells. B. Production of reactive oxygen species (ROS) was determined using MitoSOX™ assay. The bar graph represents the fold change of ROS production in cells treated with antiretroviral drugs compared to DMSO-treated control cells. C. Representative western blot image of subunits of the ETC complexes: complex V (CV-APT5A), complex III (CIII-UQCRC2), complex IV (CIV-MTCO1), complex II (CII-SDH8), and complex I (CI-NDUFB8) under DMSO and ATV treatment (5.5 and 22.5 µM). The density of the band for each complex was normalized to that of tubulin. The number below each band represents the ratio of complex expression to tubulin compared to that for DMSO-treated cells multiplied by 100; an average of at least 2 independent experiments. D. Density of the ETC protein complexes. Data (mean ±SD) were compared with DMSO treated cells and analyzed by Student’s t-test. P-values are two sided and considered significant if <0.05 (*), < 0.01 (**), or <0.001 (***).

Mitochondrial ROS production is associated with loss of mitochondrial ΔΨ ([31]). We, therefore, investigated whether the decrease in mitochondrial ΔΨ in cells treated with ATV translated to an increase in production of ROS. We determined ROS production using MitoSOX™ Red mitochondrial superoxide indicator as we previously described [23]. At days 2 and 4, cells treated with 22.5 and 30 µM ATV produced significantly higher ROS compared to cells treated with DMSO (Figure 3B).

Mitochondiral ΔΨ is generated by the proton pumps of the ETC and utilized by ATP synthase (complex V) to synthesize ATP[32]. We therefore investigated the effect of ATV on protein expression of the ETC complexes (I-V) using western blot analysis. Cells treated with DMSO and 22.5 µM ATV for 8 days were compared. Figure 3C is a representative Western blot image of subunits of ETC complexes. The density of the band for each complex was normalized to that of tubulin. The number below each band represents the ratio of complex expression to tubulin of ATV treated cells compared to DMSO treated cells multiplied by 100. Our results showed that 22.5 µM ATV treatment significantly decreased protein expression of ETC complex CI, CII, CIII, and CV (Figure 3D).

### Inhibition of NLRP3 pathway reversed mitochondrial dysfunction

We then tested mitochondrial function after ATV treatment for 6 hours with pre-incubation of AC-YVAD-CMK or MCC950. Interestingly, mitochondria ΔΨ was significantly increased after pre-incubation with either NLRP3 or caspase-1 inhibitor (Figure 4A). Furthermore, pre-incubation with either AC-YVAD-CMK or MCC950 resulted in less production of ROS (Figure 4B). These results indicated that inhibition of pyroptosis might reverse mitochondrial dysfunction to a certain extent.

**Figure 4.**
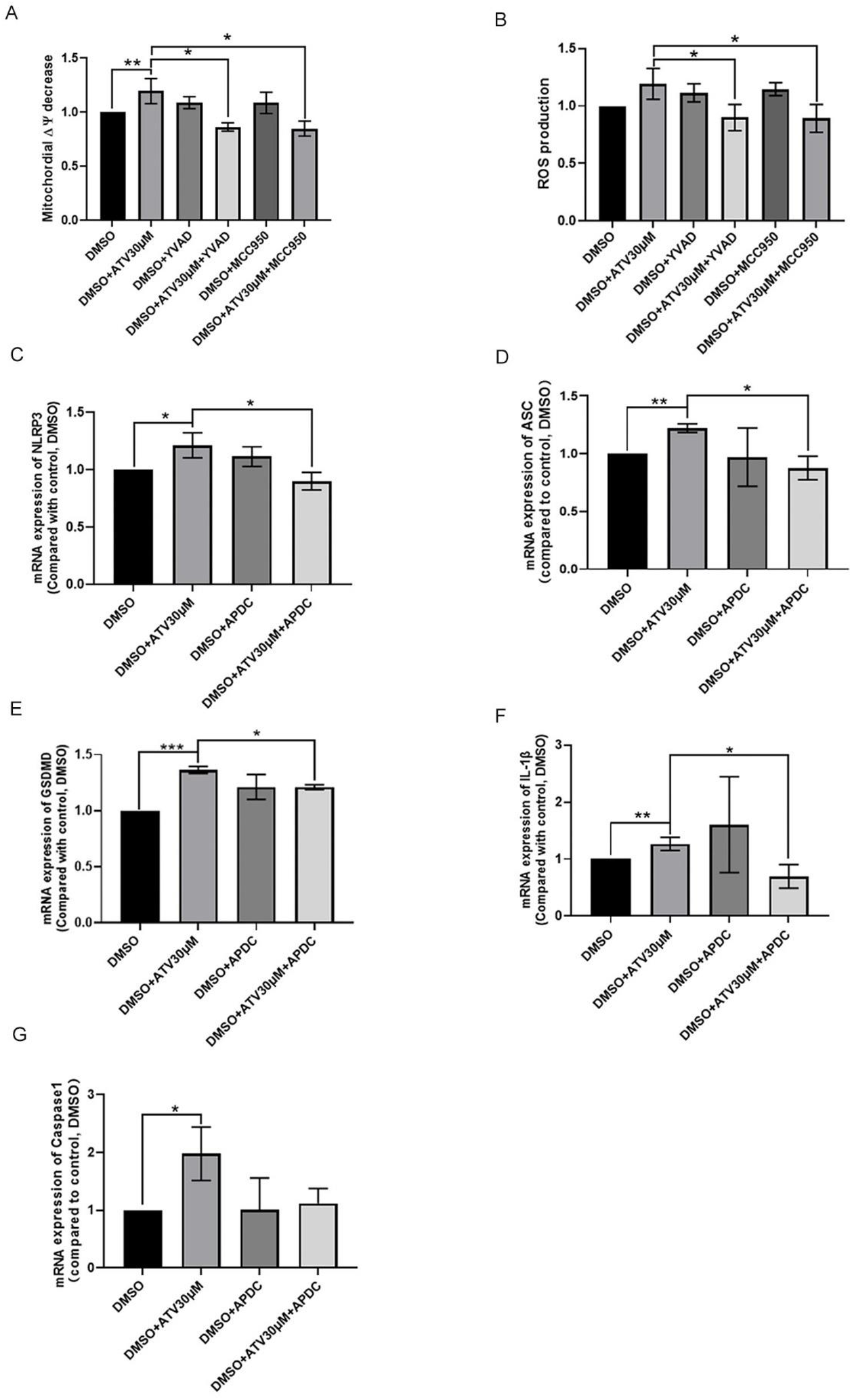
Effect of NLRP3 pathway inhibitors and reactive oxygen species (ROS) inhibitors. CEM cells were pre-incubated with AC-YVAD-AMK (20 µM) or MCC950 (1 µM) for 1 h or ammonium pyrrolidinedithiocarbamate (APDC) (2 µM) for 10 min before the following ATV treatment for 6 h. A. cells were harvested and interrogated for mitochondrial membrane potential (ΔΨ) using TMRE-ΔΨ assay Kit with flow cytometry. The bar graph represents the relative decrease of mitochondrial ΔΨ in cells treated with antiretroviral drugs compared to DMSO-treated control cells. B. Production of ROS was determined using MitoSOX™ assay. The bar graph represents the fold change in ROS production in cells treated with antiretroviral drugs compared to DMSO-treated control cells. C-G. Cells were harvested and RNA was extracted. mRNA expression level of NLRP3 (C), ASC (D), GSDMD (E), IL-1β (F) and caspase-1 (G) were determined by RT-qPCR. The bar graph represents the fold change in each parameter in cells treated with ATV compared to DMSO-treated control cells. Data (mean ±SD) were compared with DMSO treated cells and analyzed by Student’s t-test. P-values are two sided and considered significant if <0.05 (*), < 0.01 (**), or <0.001 (***).

As mitochondria and NADPH oxidase are the main source of ROS [33], we investigated the expression of key genes in NLRP3 inflammasome pathway after pre-incubation of cells with ROS scavenger ammonium pyrrolidinedithiocarbamate (APDC). Our results showed that the incubation with APDC significantly reversed the increased expression level of NLRP3, ASC, GSDMD and IL-1β in cells treated with 30 µM of ATV (Figure 4C-F). The expression level of caspase-1 decreased after pre-incubation with APDC, but didn’t reach statistical significance (Figure 4G).

## Discussion

HIV PIs are known to have concentration-dependent bi-model effect on apoptosis [5–13, 15], however, the underlying mechanism(s) are not well-understood. In this study, we sought to investigate why high concentrations of HIV PIs are pro-apoptotic. We observed that high concentrations of ATV, over and above achievable plasma concentration in patients treated with ATV, caused significant increased cell death and apoptosis in human T lymphoblastoid cell line. Further, the observed increased in apoptosis is likely due to activation of NLRP3 inflammasome and pyroptosis pathway by high concentrations of ATV. We observed an increased expression of key genes of NLRP3 inflammasome pathway, including NLRP3, caspase-1, ASC, GSDMD, IL-1β, and IL-18. Interestingly, NLRP3 and caspase-1 inhibitors reversed the pro-apoptotic effect of high concentration of ATV. Moreover, at high ATV concentrations, we observed ATV induced mitochondrial dysfunction—decreased mitochondrial ΔΨ, increased production of ROS, and decreased expressions of ETC protein complexes. Further, co-treatment of NLRP3 or caspase-1 inhibitors reversed the mitochondrial dysfunction seen with ATV treatment alone. Taken together, we surmise that high concentrations of HIV PI ATV activate NLRP3 through ATV-induced mitochondrial dysfunction. To the best of our knowledge, this is the first study to associate PI-induced apoptosis with NLRP3/pyroptosis pathway.

We found that high concentrations of ATV (15-30 µM) increased the rate of cell death and exhibited a pro-apoptotic effect in human T-lymphoblast cells. Of note, no significant changes were observed with ATV treatments at 7.5 µM or below. Similar to our findings, Kim et al. and Pyrko et al. observed that ATV had minimal or no effect at lower concentrations in KBV20C cancer cells[35] and glioblastoma cell lines[34]. Additionally, Kim et al. reported that ATV did not induce apoptosis at concentrations of 0-50 µM in human adipocytes[36]. However, other studies have reported that ATV can induce apoptosis at concentrations of 1-20 µM in human islet cells[37] and 5-50 µM in rat primary hepatocytes[38]. We believe these discrepancies across different studies are due to the varying sensitivity of the specific cell lines used *in vitro*.

The pro-apoptotic effect of high concentrations of ATV is intriguing. Similar observations were noted by Gibellini et al. and Pryko et al., who reported the pro-apoptotic impact of ATV treatment on SW872 cells at 100-200 µM [39]and on glioblastoma cell lines at 75-100 µM[34]. Several theories have been put forth to explain how high concentrations of PIs induce apoptosis, including the induction of endoplasmic reticulum stress[34, 43], activation of the intrinsic apoptosis pathway[37], and cell cycle arrest [10, 35]. We observed that four of the genes in the apoptotic pathway—caspase-1, BCL2A1, TP73, and TNFRSF1B—had over two-fold expression between DMSO and ATV treatment. Among these, the high expression of TP73 and TNFRSF1B may indicate regulation of cell cycle arrest[44, 45], while BCL2A1 upregulation may act as a protective mechanism against ROS and apoptosis[25]. The increased expression of caspase-1 is particularly noteworthy, as it plays a crucial role in the pyroptosis pathway.

Pyroptosis is an inflammatory form of programmed cell death that is accompanied by activation of inflammasomes and maturation of pro-inflammatory cytokines [28].The inflammatory reaction starts with the recognition of danger signals, including pathogen-associated molecular patterns (PAMPs) or danger-associated molecular patterns (DAMPs), by NOD-linked receptor (NLR) family, such as NLRP3-inflammasome [28]. The NLRP3 inflammasome is a multiprotein complex consisting of three components: the nucleotide-binding domain leucine-rich repeat containing (NLR) family member NLRP3, the adaptor protein ASC containing a caspase recruitment domain, and the cysteine protease caspase-1[29]. Mature caspase-1 then cleaves the caustic executor protein Gasdermin D (GSDMD), in turn to form the cell membrane pores. The mature cleaved caspase-1 also promotes the maturation of pro-inflammatory cytokines IL-1β and IL-18, which are secreted through the pores formed by GSDMD, resulting in pyroptosis [18, 19]. Pyroptosis has gained increasing attention in recent years because of its involvement in various diseases, such as tumors, inflammatory diseases, and cardiovascular diseases [19, 46, 47]. In treatment naïve people living with HIV, NLRP3 inflammasome-mediated pyroptosis is also the main pathway contributing to CD4+ T cell loss [48, 49]. Antiretroviral therapy (ART) was reported to inhibit pyroptosis based on the decreasing expression of caspase-1 in CD4+ T cells after ART [50, 51].

However, in this study, we observed that ATV, a newer generation PI, could activate NLRP3 inflammasome/pyroptosis pathway at high concentrations. Similarly, a clinical study showed that NLRP3 inflammasome activation was significantly increased in HIV infected ART-treated individuals with defective immune recovery [52]. Other classes of ART have been implicated in the activation of pyroptosis [53, 54]. Efavirenz, an NNRTI, triggered inflammation in hepatocytes and activated hepatic stellate cells (HSCs), thereby enhancing expression of NLRP3 involved inflammation [53]. Abacavir, a nucleoside reverse transcriptase inhibitor (NRTI), also induced IL-1β release through NLRP3 activation, [54]. These findings suggest that the effects and mechanism(s) of different ART drugs on activation of NLRP3 inflammasome are not class-specific and may be different.

In consideration of the fact that mitochondria is the main source of ROS, and excessive ROS production has been proposed as direct activator of NLRP3 inflammasomes [55, 56], we tested the mitochondrial function under high concentrations of ATV treatment and we observed that high concentrations of ATV could induce a decrease in ΔΨ and protein expression of ETC complex subunits CI, CII, CIII, and CV, along with an increase in ROS production, indicating mitochondrial dysfunction. Given that the activation of the NLRP3 inflammasome pathway coincided with mitochondrial dysfunction, we are intrigued by the potential relationship between these phenomena. In our study, the increased expression level of some key genes in NLRP3 inflammasome pathway activated by high concentration of ATV treatment was reversed by ROS scavenger APDC, confirming the role of excessive ROS production as activator of NLRP3 inflammasome. Interestingly, adding NLRP3 and caspase-1 inhibitors led to increased membrane ΔΨ and decreased ROS production, indicating that mitochondria function could also be rescued by inhibitors of NLRP3 pathway. This is consistent with the study of Toksoy et.al that NLRP3 depletion significantly reduced Abacavir induced mROS production [42]. Those results suggested that ROS may serve not only as a trigger, but also as an effector molecule. The relationship between mitochondrial dysfunction and the NLRP3 pathway provides valuable insights for drug discovery. The prevalence of metabolic syndrome (MetS) among people living with HIV and using antiretroviral therapy (ART) is increasing rapidly [60, 61], with mitochondrial dysfunction being a common underlying mechanism[62, 63]. Could druggable inhibitors targeting NLRP3 pathway potentially reverse mitochondrial dysfunction in HIV patients?

Although these findings are intriguing, our study has several limitations. While T cells serve as the primary target of the HIV virus, the effects observed in the CEM cell line may not fully reflect the impact on the entire human immune system. Therefore, further examination of the pro-apoptotic effect of ATV should be conducted using alternative cell lines or macrophage/dendritic cells. Additionally, we did not assess the levels of IL-1β and IL-18 in the cell supernatant. While cytokine secretion serves as compelling evidence of NLRP3 inflammasome activation, increased mRNA expression of key genes also serves as a robust indicator. Moreover, we did not investigate the regulation of cell cycle arrest under ATV treatment or how it interacts with the pyroptosis pathway. These areas merit further investigation in future studies.

In conclusion, our study illustrate that high concentrations of ATV activate NLRP3 inflammasome/pyroptosis pathway, most likely through ATV induced mitochondrial dysfunction. These results, coupled with the observed reversal effects of inflammasome inhibitors, highlight the potential of targeting NLRP3 inflammasome activity as a promising therapeutic approach for addressing ART-induced mitochondrial dysfunction in individuals living with HIV.

## Notes

**Conflicts of interest**: No conflicts of interest declared by all authors

